# *Drosophila melanogaster* is a powerful host model to study mycobacterial virulence

**DOI:** 10.1101/2022.05.12.491628

**Authors:** Esther Fuentes, Niruja Sivakumar, Linn-Karina Selvik, Marta Arch, Pere Joan Cardona, Thomas R. Ioerger, Marte Singsås Dragset

**Affiliations:** Tuberculosis Research Unit, Germans Trias i Pujol Research Institute (IGTP), Badalona, Catalonia, Spain; Comparative Medicine and Bioimage Centre of Catalonia (CMCiB), Germans Trias I Pujol Research Institute (IGTP), 08916 Badalona, Catalonia, Spain; Microbiology Department, Laboratori Clínic Metropolitana Nord, Germans Trias i Pujol University Hospital, Badalona, Catalonia, Spain; Centro de Investigación Biomédica en Red en Enfermedades Respiratorias (CIBERES), Instituto de Salud Carlos III (ISCIII), Madrid, Spain; Centre of Molecular Inflammation Research and Department (CEMIR) of Clinical and Molecular Medicine, Norwegian University of Science and Technology, Trondheim, Norway; Genetics and Microbiology Department, Universitat Autònoma de Barcelona, Bellaterra, Catalonia, Spain; Department of Computer Science and Engineering, Texas A&M University, College Station, Texas, USA

## Abstract

*Drosophila melanogaster* (*Drosophila*), the common fruit fly, is one of the most extensively studied animal models we have, with a broad, advanced, and organized research community with tools and mutants readily available at low cost. Yet, *Drosophila* has barely been exploited to understand the underlying mechanisms of mycobacterial infections, including those caused by the top-killer pathogen *Mycobacterium tuberculosis* (*Mtb*). In this study, we aimed to investigate whether *Drosophila* is a suitable host model to study mycobacterial virulence, using *Mycobacterium marinum* (*Mmar*) to model mycobacterial pathogens. First, we validated that an established mycobacterial virulence factor, EccB1 of the ESX-1 Type VII secretion system, is required for *Mmar* growth within the flies. Second, we identified *Mmar* virulence factors in *Drosophila* in a high-throughput genome-wide manner using transposon insertion sequencing (TnSeq). Of the 181 identified virulence genes, the vast majority (91%) had orthologs in *Mtb*, suggesting that the encoded virulence mechanisms may be conserved across *Mmar* and *Mtb*. Finally, we validated one of the novel *Mmar* virulence genes we identified, a putative ATP-binding protein ABC transporter encoded by *mmar_1660*, as required for full virulence during both *Drosophila* and human macrophage infection. Together, our results show that *Drosophila* is a powerful host model to study and identify novel mycobacterial virulence factors relevant to human infection.

## INTRODUCTION

Tuberculosis (TB), caused by *Mycobacterium tuberculosis* (*Mtb*), is the world’s deadliest infectious disease (ongoing SARS-CoV-2 pandemic excluded) with one death every three minutes (1). Following the World Health Organization (WHO)’s guidelines, drug-sensitive TB is treated with a combination of four antibiotics given for at least six months (1). Such a long regimen is particularly difficult in countries with socioeconomic issues. The lack of financial resources for sufficient follow-up may lead patients to finish their cures prematurely, nurturing the development of drug resistant *Mtb*, provoking even longer, more difficult treatment regimens associated with potentially severe side effects. Attractive novel strategies for improved treatments and to reduce the development of *Mtb* resistance comprise targeting host-pathogen interactions (HPIs), that is, targeting either the pathogen’s virulence by disarming it rather than targeting its viability (2), or targeting the host response factors by enhancing host protection or interfering with host factors enabling infection (called host-directed therapy) (3). Hence, we need representative animal models for dissecting HPIs. *Drosophila melanogaster* (*Drosophila*) has been proposed to be particularly fit for this purpose (4, 5).

*Drosophila*, also known as the common fruit fly, has been fundamental to our understanding of molecular mechanisms in human biology and disease, sharing 60% of its DNA with us (6). *Drosophila*’s immunity largely depends on the phagocytosis of invading pathogens by plasmatocytes (macrophage-like cells), followed by activation of the Toll or IMD (for IMmune Deficiency) pathways for antimicrobial peptide production (7). It was the identification of *Drosophila*’s Toll cascade that led to the characterization of human toll like receptors (TLRs) (8), reshaping the understanding of our own innate immune system. *Drosophila* do not possess adaptive immunity, creating the opportunity to specifically study the innate immune responses in an isolated yet *in vivo* setting. Being a work-horse in basic biological research, the *Drosophila* scientific community is both advanced and open, with state-of-the-art molecular tools and fly mutants shared at low cost. Moreover, *Drosophila* as a host model is in compliance with the 3Rs (Replace, Reduce, Refine), part of the European Union legislation Directive 2010/63/EU, for a more humane animal research. Strengthened by its short generation time and general ease to handle, the above makes *Drosophila* a powerful animal model to study innate immune response to infection.

*Mtb* infects and resides within the macrophages of the human host by (among other mechanisms) blocking acidification of phagosomes and their fusion to lysosomes (9). Likewise, Dionne et al. have shown that *M. marinum* (*Mmar*), an opportunistic human pathogen of the *Mycobacterium* genus, proliferates within *Drosophila* plasmatocytes during infection, blocking acidification of the cell’s phagosomes (10). *Mmar*, which may infect fish and extremities in humans, thrives at 29°C, as do *Drosophila*, and is thus perfectly compatible to the fruit fly in an infection setting. In fact, Dionne and others have used the *Mmar*-*Drosophila* infection model to unravel aspects of the fly’s immune responses towards mycobacterial infection (11-13). *Drosophila* has also been useful in assessing antimycobacterial drug activity, infecting the fly with either *Mmar* or *M. abcessus* (14, 15). Actually, the use of *Drosophila* in mycobacterial research was recently summarized and reviewed, emphasizing *Drosophila*’s untapped potential to study mycobacterial HPIs (4). However, to fully exploit *Drosophila* for this purpose, we need to understand which mycobacterial virulence genes and mechanisms are at play during *Drosophila* infection.

Here we show that *Drosophila* is a powerful model to study and identify mycobacterial virulence genes. By transposon insertion sequencing (TnSeq) we identified, in a genome-wide manner, 181 *Mmar* virulence genes in *Drosophila*. Interestingly, over 90% of the genes had orthologs in *Mtb*, whereas 42% of these again were identified as *Mtb* virulence genes during previously published mouse model TnSeq screening (16, 17), suggesting the *Mmar*-*Drosophila* infection model may be relevant to better understand *Mtb* virulence in mammals. Finally, we knocked out one of the identified novel virulence genes, *mmar_1660* (a conserved putative ATP-binding protein of an ATP-binding cassette (ABC) transporter), and demonstrated that the mutant was attenuated for growth within *Drosophila* as well as human macrophages.

## RESULTS

### The established mycobacterial virulence factor EccB1 is required for *Mmar* virulence in *Drosophila*

We wanted to investigate whether *Drosophila* is a suitable host model to study mycobacterial virulence and hypothesized that, if so, established mycobacterial virulence factors should be required for *Mmar* infection in *Drosophila*. Hence, we infected *Drosophila* with an *Mmar* mutant in EccB1. EccB1 (encoded by *eccB1*) is a core component of the 6 kDa early secretory antigenic target (ESAT6) Type VII secretion system 1 (ESX-1) (18). ESX-1 secretion enables transport of specific protein substrates involved in *Mtb* virulence, like CFP-10 and ESAT-6, across the complex mycobacterial cell wall, and is considered a hallmark in mycobacterial virulence (19). Indeed, a mutation in *Mmar eccB1* (transposon insertion mutation; *eccB1::tn*) increased the survival of the flies upon infection, compared to *Mmar* wild type (wt) infection (Fig. 1A). Complementary, the *eccB1* mutant showed attenuated growth within the flies (Fig. 1B), most probably explaining the prolonged survival of the mutant-infected flies. Hence, we observed the expected virulence-impaired phenotype (attenuated growth within the host and prolonged host survival) of an established mycobacterial virulence gene mutant, suggesting that *Drosophila* is indeed a suitable host model to study mycobacterial virulence.

**Figure 1.**
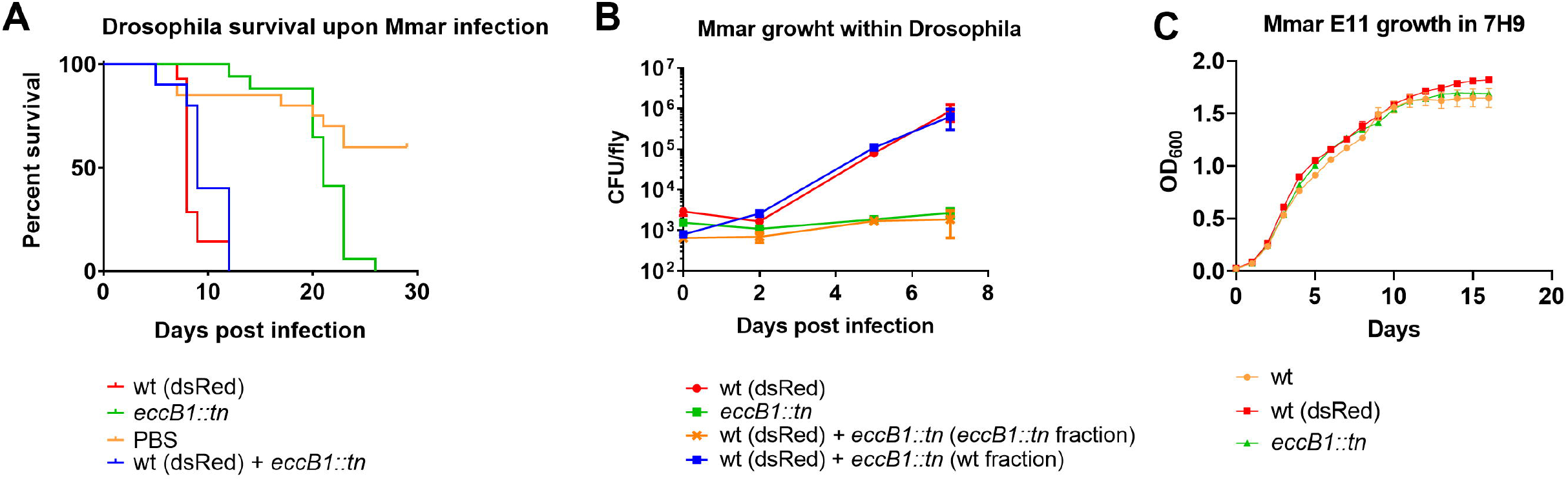
*Drosophila* infected with 5000 CFU/fly of *Mmar* E11 strain wt (dsRed), *eccB1::tn*, a 1:1 mix of wt (dsRed) or *eccB1::tn*, or PBS. Fly survival (A) and CFU per fly (A) was recorded over the course of infection. For CFU measurements, wt (dsRed) and *eccB1::tn* were selected on hygromycin or kanamycin, respectively, and data represent means ± standard error of the mean (SEM) for three individual flies per condition and time point. For fly survival, data represent the percent survival of initial 30 flies per condition and were recorded daily. (C) *Mmar* E11 wt, wt (dsRed), and *eccB1::tn* growth *in vitro* (7H9 medium) as measured by OD_600_. Data represent means ± SEM of three technical replica recorded every 24 hours.

### *Drosophila* is a suitable host model to screen for novel mycobacterial virulence genes

Next, we asked whether *Drosophila* is suitable to identify *novel* mycobacterial virulence genes. We and others have, by TnSeq, previously identified mycobacterial virulence genes in a genome-wide manner using mouse (16, 17, 20, 21), cattle (22), or ameba (23, 24) as host infection models. Discovery of virulence genes by TnSeq is based on infecting the host with a bacterial high-density transposon (tn) mutant library, followed by harvesting the library for analysis after infection, comparing its mutant constituents to the input library using massive parallel sequencing (25). Mutants that are growth-impaired within the host, but not on standard agar medium, are considered virulence genes. Hypothetically, though, a mutant deficient in for instance ESX-1 secretion could be rescued by mutants sufficient in ESX-1 secretion residing within the same plasmatocyte. To investigate whether this could obscure our screen for virulence genes when we pass a high number of mutants through one single fly, we infected *Drosophila* with 5000 colony forming units (CFU)/fly of either *Mmar* wt, *eccB1::tn* mutant or a 1:1 mix of *Mmar* wt and *eccB1::tn* mutant. By using an *Mmar* wt strain carrying a plasmid conferring hygromycin resistance and dsRed expression, wt (dsRed), in combination with the *eccB::tn* mutant carrying kanamycin resistance encoded within the tn insert, we could by antibiotic selection separate the growth of the mutant from wt growth during co-infection. As seen in Figure 1B, the *eccB1* mutant was still growth-attenuated within the flies when co-infected with *Mmar* wt, while the co-infected flies died similarly to those infected by wt only (Fig. 1A). The strains grew comparably in 7H9 liquid medium, and the wt (dsRed) strain grew similarly to the wt strain (Fig. 1C). From this we take that virulence mutants are not rescued by co-infection with other mutants intact in their version of the virulence gene in question, although this may vary depending on the virulence mechanism. Finally, by day seven post infection the wt bacteria had doubled approximately seven times, providing room to differentiate between mutants in virulence and non-virulence genes during TnSeq screening (Fig. 1A). Taken together, our results suggest that *Drosophila* is a suitable host model to screen for novel mycobacterial virulence.

### Genome-wide identification of *Mmar* virulence genes in *Drosophila*

We aimed to identify novel Mmar virulence genes in *Drosophila*. Hence, we constructed, using ϕMycoMarT7 (26), and sequenced a high-density transposon insertion library in the E11 strain. The obtained library contained mutants in 80% of TA sites covering 96.5% of the genes. Moreover, we uncovered 430 essential (E) genes, 130 genes conferring growth defect (GD) when disrupted, 4314 non-essential (NE) genes, and 79 genes conferring growth-advantage (GA) when disrupted (S1 Datasets A). Combining ES and GD categories, the 560 genes that are essential or cause a growth-defect when disrupted in are line with what has been observed in *Mtb* (27, 28). In a previous TnSeq study of *Mmar* (24), 300 genes were identified as being essential *in vitro*, of which 82% (247) are also in the ES or GD category in our data (S1 Datasets A).

To specifically identify *Mmar* virulence genes during *Drosophila* infection, we passed the generated library through *Drosophila* (250 flies, 5000 CFU/fly). We subjected the input and output libraries to TnSeq (the latter harvested from the flies seven days post infection) and found 181 genes that were required for optimal growth within the flies, based on a permutation test (“resampling analysis” (29)) of the difference in mean tn insertion counts per gene in libraries that had undergone fly infection (output) versus libraries grown under *in vitro* condition (input) (log fold change <0 and adjusted P-value <0.05) (S1 Datasets B). Among the virulence genes identified were those encoding established mycobacterial virulence factors, like phthiocerol dimycoceroserate (PDIM, a cell wall lipid, *mmar_1667, _1770-1771*), components of the ESX-1 secretion system (10 genes genes between *mmar_5399-5459*), and the LytR-CpsA-Psr domain-containing protein CpsA (*mmar_4966*), but also novel genes never before associated with virulence. We also identified genes (15) that conferred *Mmar* growth advantage within *Drosophila* when disrupted (log fold change >0 and adjusted P value <0.05), making them the mere opposite of virulence genes (S1 Datasets B). An example of these are the *eccA1* (*mmar_5443*). Our result taken together, we identified 181 mycobacterial virulence factors during *Mmar* infection of *Drosophila*.

### The relevance of *Mmar* virulence in *Drosophila* to *Mtb* infection

To investigate the potential relevance the *Mmar* virulence genes in *Drosophila* may have to *Mtb* infection, we compared them to the entire gene pool of *Mtb*. We found that most of them (90%) had orthologs in *Mtb* (Fig. 2). Moreover, when we compared the *Mmar* virulence genes to those identified previously during *Mtb* mouse infection (16, 17), 42 % of them had *Mtb* orthologs that were also required for virulence in mice. The 88 *Mmar* virulence genes that did not overlap with previous screening for *Mtb* mouse model virulence genes, may represent orthologs of novel *Mtb* virulence genes. Together, our findings suggest that mechanisms of the identified virulence genes may be conserved across *Mmar* and *Mtb* and across host model species. The relevance of the genes identified with the *Mmar*-*Drosophila* infection model may therefore translate to *Mtb* infection.

**Figure 2.**
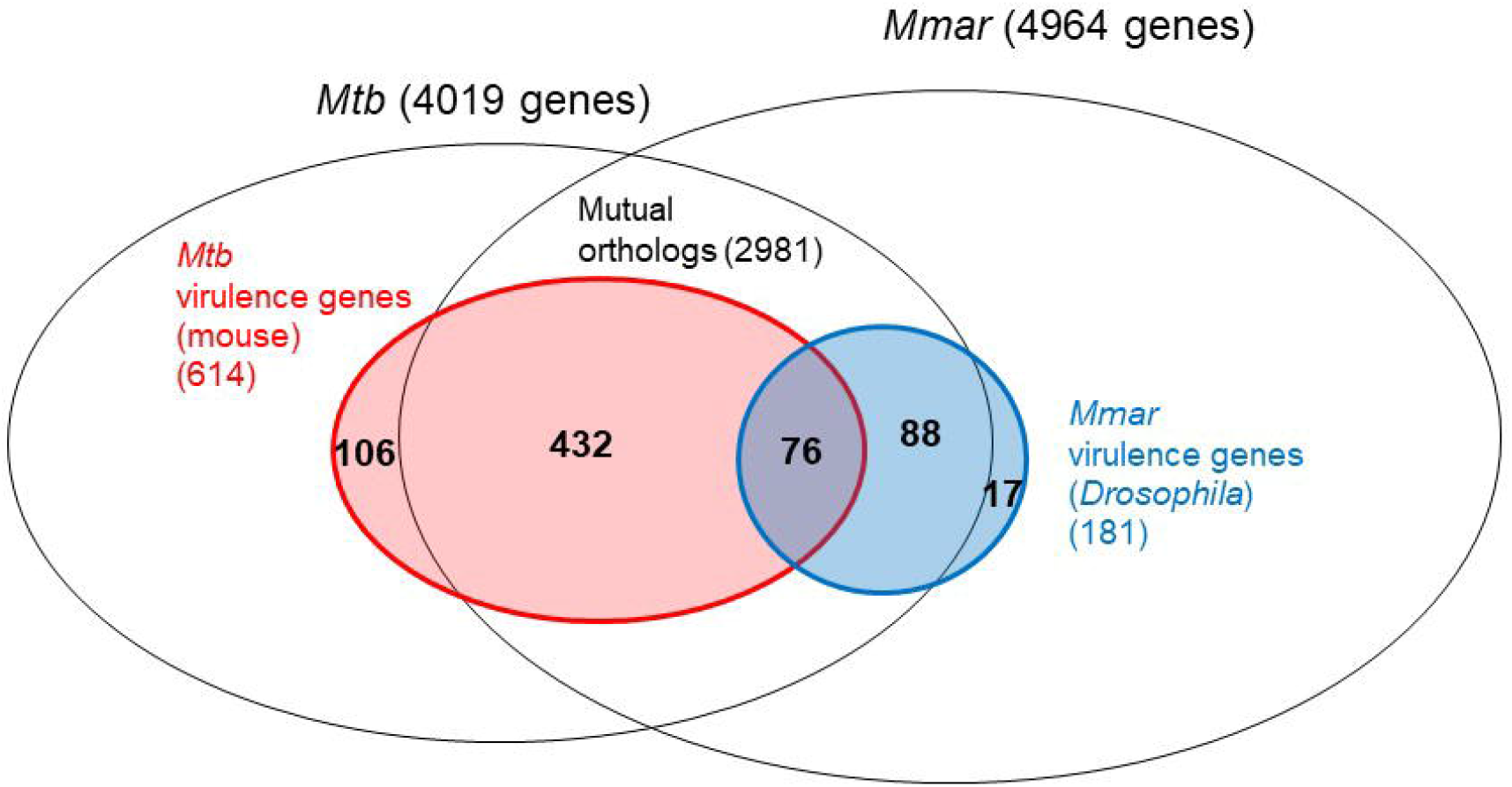
Venn diagram illustrating *Mmar* virulence genes in *Drosophila* (blue) and *Mtb* virulence genes in mice (red) (as defined by Zhang et al. and Smith et al. combined (16, 17)) overlayed the entire pool of *Mmar* (E11 strain) and *Mtb* (H37Rv strain) genes and their mutual orthologs.

### *Mmar_1660* is a novel mycobacterial virulence gene

To validate our screen, we created targeted knock out mutation of one of the novel virulence genes we identified, *mmar_1660*, ortholog of *Mtb rv3041c*. This gene encodes a putative conserved ATP-binding protein ABC transporter, MMAR_1660 (30). When we infected *Drosophila* with *Mmar* wt, *Mmar* Δ1660, and *Mmar* Δ1660 complemented strains, Δ1660-infected flies survived on average one day longer than wt-infected (Fig. 3A). Complementing the knockout mutant with expression of the intact *mmar_1660* gene from a genomic integration site, partially restored the attenuated phenotype of the Δ1660 mutant (Fig. 3A). All strains grew comparably *in vitro* in standard liquid medium (Fig. 3B). These results validate *mmar_1660* as a virulence gene during *Drosophila* infection.

**Figure 3.**
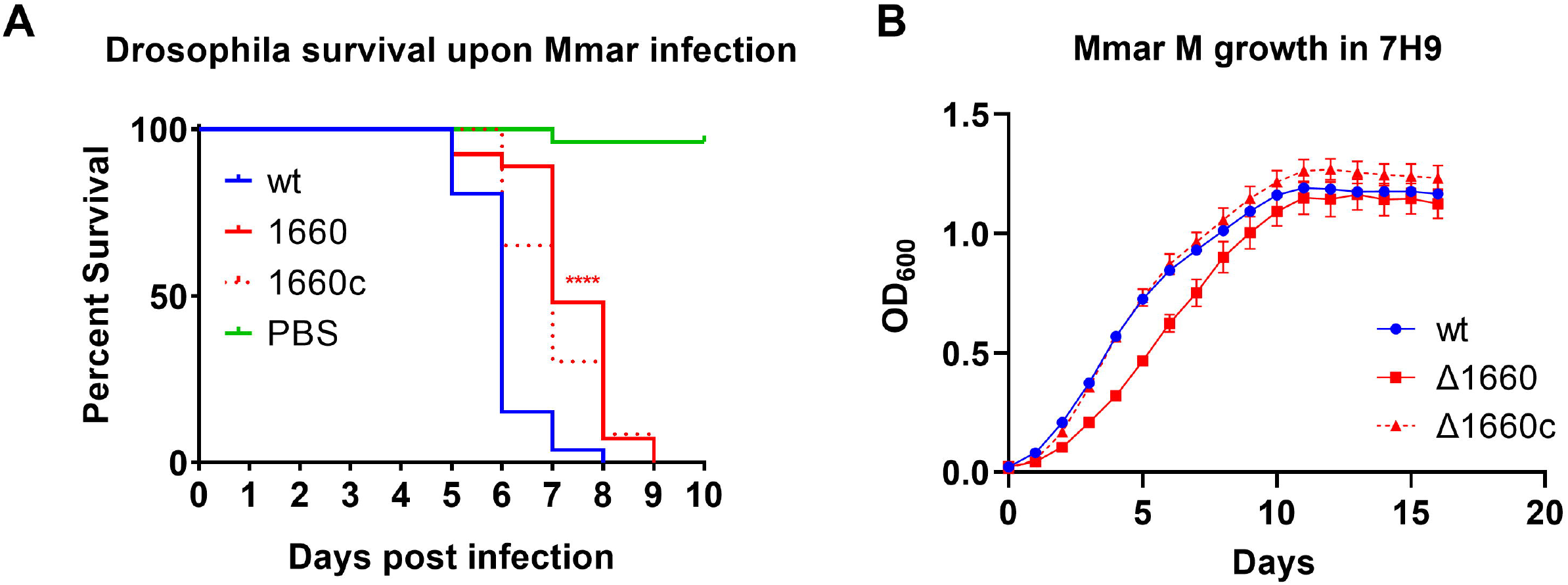
(A) *Drosophila* infected with 5000 CFU/fly of *Mmar* M strain wt, Δ1660, Δ1660 complemented (Δ1660c), or PBS, with their survival recorded over the course of infection. The four red asterisks represent a statistically significant difference (P value <0.0001) between wt and Δ1660-infected flies as calculated by Log-rank Mantel-Cox testing using GraphPad Prism 9. Data represent the percent survival of initial 30 flies per condition and were recorded daily. (B) *Mmar* M wt, Δ1660, Δ1660 complemented (Δ1660c) growth *in vitro* (7H9 medium). Data represent means ± SEM of three technical replica recorded every 24 hours.

Further, we wanted to investigate whether *mmar_1660* contributed to virulence in human macrophages. Therefore, we infected human induced pluripotent stem cells (iPSC)-derived macrophages with the *Mmar* wt, *Mmar* Δ1660, and the *Mmar* Δ1660 complemented strains. The macrophages survived longer when infected with the Δ1660 mutant compared to wt, as evident from measuring 1) tryphan blue staining of dead macrophages (Fig. 4A), 2) lactate dehydrogenase (LDH) release from dead macrophages (Fig. 4B), 3) microscopic visualization of infected macrophages (Fig. S1), and 4) indirectly by the number of *Mmar* CFU released into the cell culture medium over the course of infection (Fig. 4C). Complementary, the intracellular growth of *Mmar* Δ1660 was impaired compared to wt (Fig. 4D). We saw a partial rescue of the Δ1660-mediated virulence-impaired phenotype by the complemented strain (Fig. 4B and S1). In summary, we show that *mmar_1660* is required for full virulence in *Drosophila* as well as in human macrophages.

**Figure 4.**
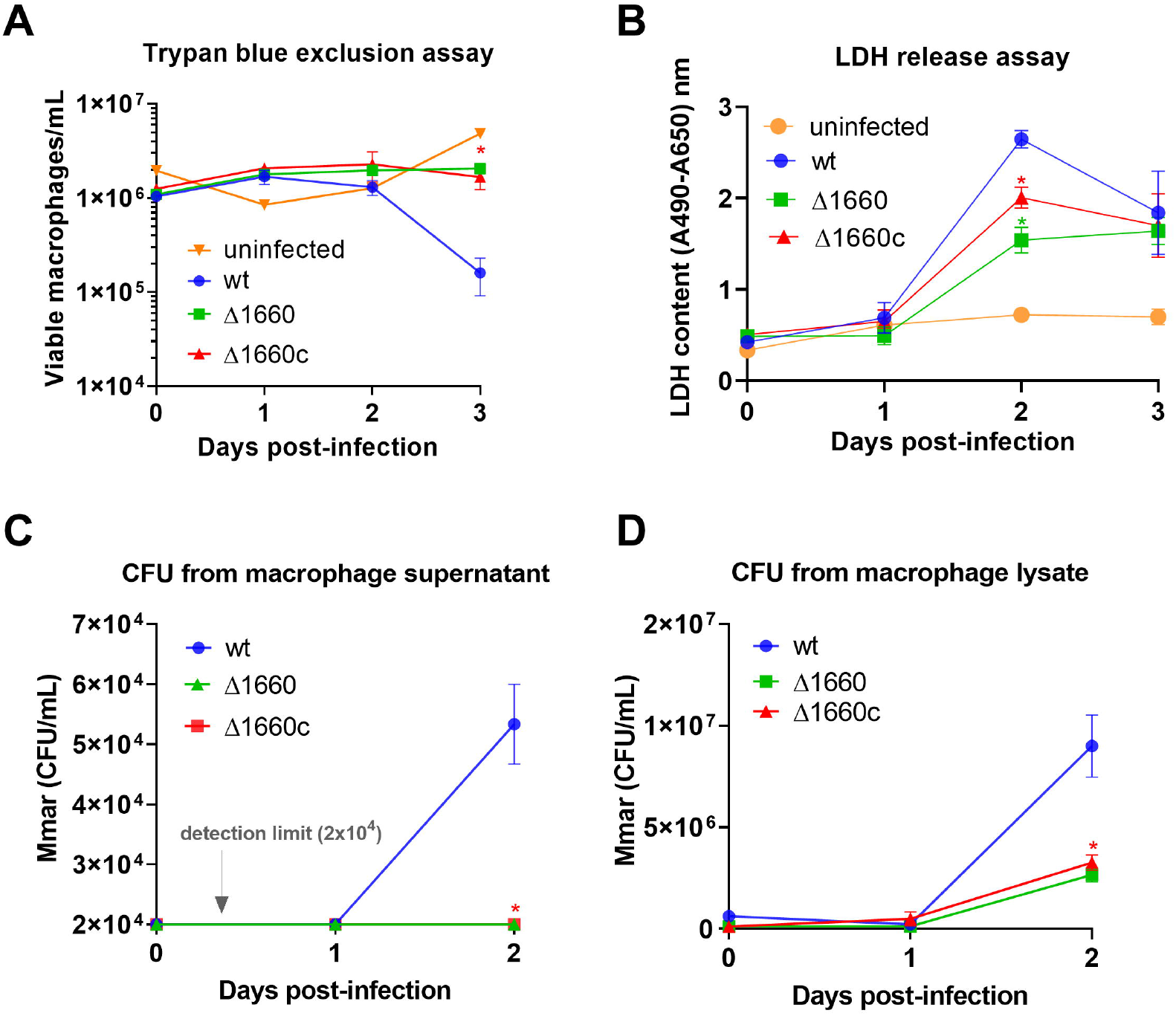
iPSC-derived human macrophages were infected with *Mmar* M strain wt, Δ1660, or Δ1660 complemented (Δ1660c). (A) Macrophage viability was determined by trypan blue exclusion assay. Data represent means ± SEM of three technical replica per condition. The red asterisk represents a statistically significant difference (P value <0.05) between wt and Δ1660-infected macrophages as calculated by Mann-Whitney U test (one-tailed) using GraphPad Prism 9. (B) *Mmar*-induced macrophage cytotoxicity was determined by absorbance measurements of LDH release, where the detection and reference wavelengths were set at 450 and 650 nm, respectively. Data represent means ± SEM of three technical replica per condition. The red and the green asterisks represent a statistically significant difference (P value <0.05) between wt and Δ1660-infected macrophages and between Δ1660 and Δ1660c-infected macrophages, respectively, as calculated by Mann-Whitney U test (one-tailed) using GraphPad Prism 9. (C, D) Bacterial burden was assessed by measuring CFU from supernatant (C) and lysed cells (D). Data represents means ± SEM from three technical replicates per condition. The red asterisk represents a statistically significant difference (P value <0.05) between wt and Δ1660-infected macrophages calculated by Mann-Whitney U test (one-tailed) using GraphPad Prism 9. For CFU in the supernatant (C), the statistical difference was calculated using the detection limit value for Δ1660-infected macrophages, from which data were below the detection limit.

## DISCUSSION

We hypothesized that *Drosophila* could be a powerful future host model to study mycobacterial virulence, due to its 1) short generation time, 2) general ease to handle, 3) state-of-the-art molecular tools and fly mutants readily available at low cost, and 4) compliance with the 3Rs for a more humane animal research. *Drosophila* has been fundamental to our understanding of mechanisms underlying human biology, the response to infection included (7, 8). While *Drosophila* has previously shown useful to understand the host’s responses towards mycobacterial infections (4), we show here that it is a powerful host model to study mycobacterial virulence from the pathogen’s perspective as well, making it particularly apt to unravel mechanisms of HPIs relevant to pathogenic mycobacteria.

By TnSeq of *Mmar* tn libraries before and after *Drosophila* infection, we found 181 genes that were required for full *Mmar* virulence. Of these, many were already established mycobacterial virulence genes, while others were novel to the field (42 and 58% respectively, based on *Mtb* mouse model TnSeq virulence screening). 91% of the identified *Mmar* virulence genes had orthologs in *Mtb*, likely reflecting that virulence mechanisms relevant to innate immunity and growth within the macrophage/plasmatocyte are conserved across the pathogenic mycobacterial *Mmar* and *Mtb*, and across arthropod (*Drosophila*) and vertebrate (mouse) host models.

Among the already established mycobacterial virulence genes we identified during the *Mmar*-*Drosophila* infection screening, were genes encoding factors involved in macrophage intracellular survival and escape, affirming that *Drosophila* is apt to study virulence associated to overcoming innate immunity. For instance, PDIM is thought to contribute to phagosomal escape and macrophage exit (31), the ESX-1 secretion system is thought to secrete immune modulating effectors into the phagosome and cytosol in addition to facilitate phagosomal escape (32), while CpsA contributes to evade macrophage killing by inhibiting the lysosomal-trafficking pathway LC3-associated phagocytosis (33).

We validated *mmar_1660* (the ortholog of *Mtb rv3041c*) as required for full virulence during *Drosophila* and human macrophage infection. This gene encodes a putative ABC transporter, predicted to be involved in active transport of possibly iron across the membrane (30). Iron is an essential nutrient for most organisms, including mycobacteria which rely on various strategies to acquire this precious metal during infection (34), like siderophore and hemophore production to scavenge ferric iron and heme, respectively (35). *mmar_1660* may therefore be involved in a novel mechanism to obtain iron during infection, or it may encode a hitherto unknown component of already known iron uptake mechanisms. In fact, *Mmar* orthologs of genes known to be involved in *Mtb* siderophore uptake (*irtA, irtB*) and biosynthesis (*mbtD, mbtG*) (36, 37), were detected as virulence genes in our screen, demonstrating that mycobacterial genes with a function in iron uptake may indeed be discovered using *Drosophila*.

Among the *Mmar* genes that mediated growth advantage within *Drosophila* when disrupted, was *eccA1* of the ESX-1 secretion system. While several ESX-1 genes were required for full *Mmar* virulence, the presence *eccA1* seem to rather slows down *Mmar* proliferation within the flies. EccA1 is thought to regulate mycolic acid lipid synthesis (38), in addition to facilitate ESX-1-mediated secretion of the key mycobacterial virulence factors ESAT-6 and CFP-10 (39). Actually, Weerdenburg et al. found that, on a cell culture level, EccA1 was required for virulence in mammalian but not in protozoan cells (24). EccA1 may therefore be involved in fine-tuning ESX-1 secretion, and in certain host species its absence may lead to *Mmar* growth advantage, perhaps by removing a functional brake for secretion.

## CONCLUDING REMARKS

Here, for the first time, we show that *Drosophila* is a powerful model to study and identify mycobacterial virulence genes in a manner that is relevant to infection of human cells, complementing its role as a model to study innate immune responses towards mycobacterial infections. Future applications of the *Mmar*-*Drosophila* infection model comprise dissecting HPIs, for instance by pinpoint specific genetic interactions between *Mmar* and *Drosophila* by TnSeq of *Mmar*-infected *Drosophila* host response mutants.

## MATERIALS AND METHODS

### Bacterial strains and growth conditions

The *Mycobacterium marinum* (*Mmar*) wt strains used in this study were *Mmar* E11 and *Mmar* M (NCBI GenBank accession numbers CP000854.1 and HG917972.2, respectively). The wt strains, in addition to the mutant strains *Mmar* E11 *eccB1::tn* (kanamycin resistant, (24)) and *Mmar* E11 dsRED (pSMT3dsRed, hygromycin resistant), were kind gifts from Wilbert Bitter at Vrije Universiteit Amsterdam. The construction of *Mmar* M Δ1660 and Δ1660 complemented strains are explained further below. *Mmar* strains were cultured in Middlebrook 7H9 (BD Difco) supplemented with 0.2% glycerol, 0.05% Tween 80 and 10% ADC (50 g BSA fraction V, 20 g dextrose, 8.5 g NaCl, 0.03 g catalase, dH2O up to 1 L) for liquid growth, with 20 μg/ml kanamycin or 50 μg/ml hygromycin added where required. For solid growth, Middlebrook 7H10 (BD Difco) supplemented with 0.5% glycerol and 10% ADC was used, with 20 μg/ml kanamycin or 50 μg/ml hygromycin added where required. *In vitro* growth curves were performed as previously described for *M. avium* (20), using the Bioscreen growth curve reader (Oy Growth Curves Ab Ltd.).

### *Mmar_1660* knockout and complemented strains

To create the *mmar_1660* knockout strain (Δ1660), bacteriophage-mediated allelic exchange using pYUB1471 and phAE159 was used (kind gift from William R Jacobs at Albert Einstein College of Medicine), following published protocols (40). In short, around 1000 bp upstream and downstream of *mmar_1660* were amplified, and cloned into pYUB1471, creating pYUB_1660. pYUB_1660 was further cloned into phAE159, which was used to transduce *Mmar* M to induce allelic exchange in order to replace the *mmar_1660* gene with genes encoding hygromycin resistance (hygR) and levansucrase (sacB) genes. Positive clones were selected on 7H10 agar containing hygromycin and further validated by PCR. The Δ1660 complemented strain was created by cloning *mmar_1660* into pFLAG_attP_kanR by Gibson Assembly (New England Biolabs) and co-transformed into *Mmar* M wt together with pMA_int (41). pFLAG_attP_kanR was created by replacing the hygR gene from pFLAG_attP (41) with the kanamycin gene from pMSP12-cfp (kind gift from Christine Cosma and Lalita Ramakrishnan (42)). pMA_int and pFLAG_attP were kind gifts from Markus Seeger (Addgene plasmids #110096 and #110095, respectively (41)).

### *Mmar* transposon mutant library

The high-density transposon mutant *Mmar* E11 library was prepared as previously described for *M. avium*, using ϕMycoMarT7 (kind gift from Eric Rubin at Harvard T.H. Chan School of Public Health (26)) (20), except for heating medium and bacterial cultures to 30°C as opposed to 37°C prior to transduction, and incubating the library at 30°C as opposed to 37°C on 7H10 plates with Tween 80 (0.05%) and kanamycin (20 μg/ml) for 2 weeks.

### Bacterial infection stocks

*Mmar* strains and tn library were prepared to obtain single-cell suspensions prior to *Drosophila* and human macrophage infections. Strains were cultured to stationary phase, pelleted down by centrifugation, and resuspended in 7H9 medium with 0.2% Tween 80 to resolve large clumps. Cultures were then pelleted again and resuspended in 7H9 media with 15% glycerol. The resuspension sat at room temperature for 30 minutes to let clumps fall to the bottom before the supernatant was transferred and aliquoted to cryotubes for storage at -80°C. To calculate CFU/mL of the stocks, we either measured OD_600_ and calculated with OD_600_ of 1 equating to 3.4 × 10^7^ CFU/mL (for macrophage infection) or diluted and spotted the stocks on 7H10 agar plates to determine the CFU/mL (for *Drosophila* infection).

### *Drosophila* strains

The *Drosophila melanogaster* (*Drosophila*) strain used for TnSeq was a cross between the Transgenic RNAi Project (TRiP) control fly AttP40 (y v; attP40, y+, stock #36304 at the Bloomington *Drosophila* Stock Center, BDSC) and a tubulin-Gal4 driver fly. For all other *Drosophila* infections the Oregon R-C (Flybase ID FBsn0000276, stock #5 at the BDSC) was used. Flies were bread at 25ºC with constant light:dark cycles of 12 hours each and a humidity of 70%. The Bloomington standard cornmeal formulation containing yellow cornmeal, corn syrup solids, inactive nutritional yeast, agar and soy flour was used to feed flies (pre-mixed dry version available at Genesee Scientific).

### *Drosophila* infection, survival and CFU assay

Infections were performed in 3 to 5-days-old flies. Flies were infected using a Nanoject II (Drummond Scientific Company) set to inject 13.8 nl, and glass needles prepared using a PB-7 needle puller (NARISHIGE) and were not exposed to CO_2_ anesthesia for more than 15 minutes during the process. Bacterial infection stocks were diluted to 5000 CFU/13.8 nl in 1:1 ratio of phosphate-buffered saline (PBS) to Brilliant Blue (Sigma Aldrich) and flies were incubated at 29°C after infection. For CFU assays, flies were put briefly into 70% ethanol before washed once with PBS and homogenized in 100 μl PBS using a pestle. The homogenates were spotted in a 10-fold dilution series on 7H10 agar plates containing 20 μg/ml kanamycin and 1.25 μg/mL Amphotericin B (to eliminate the fly’s microbiota). For each CFU count, three flies per condition were harvested, homogenized and treated separately during dilutions and spotting. Two technical replica per fly were spotted. The agar plates were grown for around 10 days at 30°C before counting CFUs. For TnSeq, 250 flies were infected with 5000 CFU of *Mmar* E11 tn library per fly in three biological replica (3 × 250 infected flies). 10 and 10 flies were put briefly into 70% ethanol before washed once with PBS and homogenized in 200 ul PBS using a pestle. The homogenates from 10 and 10 flies were plated onto 15 cm diameter 7H10 agar plates containing 20 μg/ml kanamycin, 1.25 μg/mL Amphotericin B and 0.05% Tween 80, ending up with around 25 plates per library, and grown for 14 days at 30°C. The three biological replica were treated separately throughout the experiment. For survival assays, 30 flies were infected per condition and the number of dead and live flies was noted every morning.

### Transposon insertion sequencing, TnSeq

The transposon library was harvested and pooled by scraping agar plates with colonies. Total DNA was purified using Masterpure DNA purification kit (Epicentre) and prepared for TnSeq by PCR amplification of transposon-genome junctions and adapter ligation as previously described (43). The samples were sequenced on an Illumina NextSeq 2000, generating around 12-15 million 150+150 bp paired-end reads per sample.

### Bioinformatic analysis of TnSeq datasets

The reads were processed using TPP in TRANSIT (44), which counts reads mapping to each TA dinucleotide site. Beta-Geometric correction was applied to the datasets to adjust for skewness (45). Essential genes were identified using a hidden Markov model (HMM), incorporated into TRANSIT (44), as in described in more detail previously for *M. avium* (20). Virulence genes (comparative analysis between input and output tn libraries; determining statistical differences in sum of tn insertion counts in genes within library selected *in vitro* versus after infection) were identified using the “resampling” algorithm incorporated into TRANSIT (44).

### Macrophage infection and CFU assay

Human induced pluripotent stem cell (iPSC) were obtained from European Bank for induced pluripotent Stem Cells (EBiSC, https://ebisc.org/about/bank), distributed by the European Cell Culture Collection of Public Health England (Department of Health, UK) and produced into monocytes as previously described (46). Monocytes were seeded in 96-well plates and differentiated into macrophages in RPMI 1640 with 10% fetal calf serum (FCS) and 100 ng/mL M-CSF (Prepotech, 300-25). At day six of differentiation, the cells’ medium was changed to RPMI 1640 with 10% FCS and the respective *Mmar* strains at a multiplicity of infection of 1:2 *Mmar* CFU per macrophage followed by incubation at 30°C and 5% CO_2_. Uninfected cells were included as controls. After two hours incubation, the cells were washed once with phosphate-buffered saline (PBS) to remove extracellular bacteria before adding RPMI 1640 with 10% FCS. At 0, 1 and 2 days post infection supernatant was collected and cells were washed once again with PBS before being lysed in PBS with 0.5% Triton X-100 (Sigma-Aldrich). The supernatant and cell lysate were then spotted in a 10-fold dilution series on 7H10 agar plates containing 1.25 ug/mL Amphotericin B (ThermoFisher Scientific, to avoid possible fungal infections). After one week incubation at 30°C the plates were taken out to count CFUs from dilutions with optimal countable range.

### Trypan Blue Exclusion assay

Trypan blue exclusion assay was performed 0, 1, 2 and 3 days post infection to assess cell viability. First, supernatant from infected macrophages was removed and cells were washed once with PBS followed by Accutase (Sigma-Aldrich) treatment for 1 hour at 37°C to detach adherent cells. As a final detachment step, the bottoms of the wells were scraped in circular motion with a sterile pipette tip. The cell suspension was then carefully mixed with 0.4% Trypan Blue (GE Healthcare) in a 1:1 ratio and loaded onto cell counting slides (NanoEnTek). The slides were immediacy read with Countess™ 3 Automated Cell Counter (Invitrogen) for viability with optimal range set between 1 × 10^5^ and 4 × 10^6^ cells/mL according to manufacturer’s protocol.

### LDH release assay

Lactate dehydrogenase (LDH) release from infected macrophages were determined to assess cytotoxicity 0, 1, 2 and 3 days post infection. Supernatant from infected cells were harvested, pelleted down to remove cellular debris and subjected to colorimetric analysis with LDH Cytotoxicity Assay kit (Invitrogen) according to manufacturer’s protocol. Absorbance values for LDH activity were read at detection wavelength 450 nm and reference wavelength 650 nm using iMark™ Microplate Absorbance Reader (Bio-Rad).

## ACKNOWLEDGEMENTS

We thank Claire Louet for technical assistance during the macrophage infection experiments and Prof. Trude Helen Flo and Dr. Marit Bugge for providing iPSC-derived macrophages, all at Norwegian University of Science and Technology. This research was funded by the Research Council of Norway (www.forskningsradet.no) through its Centers of Excellence funding scheme project number 223255/F50 (CEMIR), and through project number 249901 (MSD), by a Research Grant (2020) from the European Society of Clinical Microbiology and Infectious Diseases (ESCMID) to (MSD), by “La Caixa” Foundation (ID 100010434), under agreement LCF/PR/GN16/10290002, by Spanish Government-FEDER Funds through PI17/01511 grant, and by the “CIBER Enfermedades Respiratorias” Network (CIBERES).

## SUPPLEMENTARY MATERIAL

**S1 Figure** Visualization of iPSC-derived macrophages either uninfected or infected with *Mmar* M strain wt, Δ1660, or Δ1660 complemented (Δ1660c) 0 and 3 days post infection, using EVOS™ FL Auto 2 Imaging System (Invitrogen) with autofocus set at first field each area (each area was defined as one well in a 96-well plate), transmitted light channel and ×10 objective.

**S1 Datasets**. (A) Essential gene analysis: *Mmar* E11 *in vitro* genetic requirement (determined by HMM in TRANSIT (44)). (B) Virulence gene analysis: *Mmar* E11 genes required for infection in *Drosophila* (determined by resampling analysis in TRANSIT (44)). Virulence genes = log fold change >0 and adjusted P-value <0.05.

