## Supplementary figures and images for "*Drosophila melanogaster* is a powerful host model to study mycobacterial virulence"

### S1 Figure

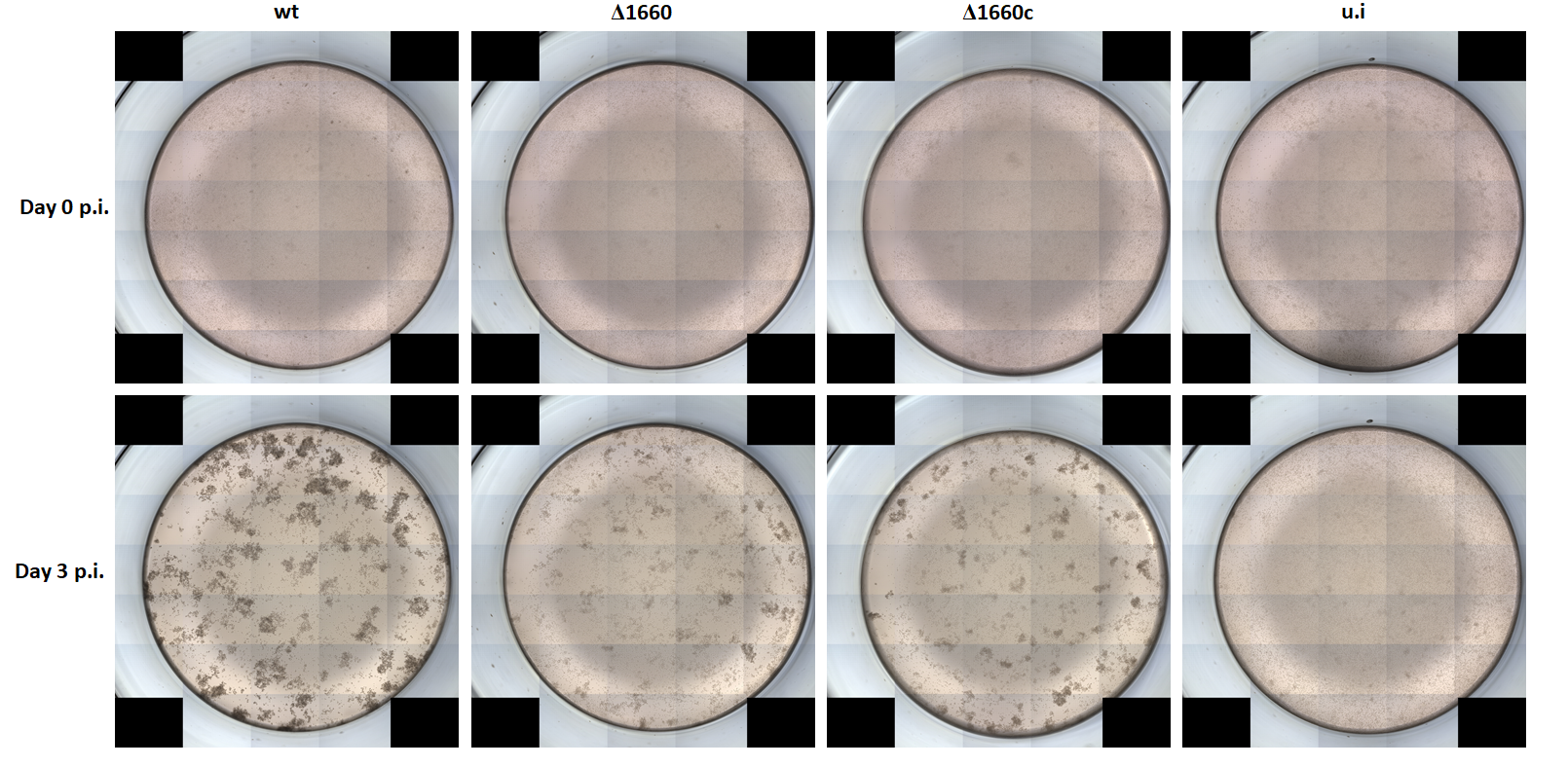
